# Larger is not better: No mate preference by European Common Frog (*Rana temporaria*) males

**DOI:** 10.1101/2021.05.28.446140

**Authors:** Carolin Dittrich, Mark-Oliver Rödel

**Author notes:** E–mail corresponding author.

## Abstract

According to classical sexual selection theory, females are the choosy sex in most species. Choosiness is defined as the individual effort to invest energy and time to assess potential mates. In explosive breeding anurans, high intrasexual competition between males leads to a sexual coercion ruled mating system, where males could have evolved preferences for specific female traits. In the current study, we tested male mating preference in the explosive breeding European Common Frog without intrasexual competition. We hypothesized that males show preferences towards larger female body size in the absence of male competition. We conducted mate choice experiments, placing a male and two differently sized females in a box and recorded their mating behavior. Males did not show any preference considering female body size, neither in the attempt to grab a female nor during the formation of pairs. We witnessed a high failure rate of male mating attempts, which might make the evolution of mate choice too costly. However, small males are faster in attempting females, which could be an alternative strategy to get access to females, because their larger competitors have an advantage during scramble competition. Nonetheless, in successfully formed pairs, the females were on average larger than the males, an observation which deviated from our null-model where pairs should be of similar size if mating would be random. This indicates that selection takes place, independent from male mating preference or scramble competition.

## Introduction

Research on sexual selection, exploring the mechanisms that lead to female/male mate choice and the evolution of different mating systems that facilitate non-random mating, has increased considerably in recent years (Janetos 1980; Ryan and Keddy-Hector 1992; Paul 2002; Edward and Chapman 2011). Studies addressing the theory of sexual selection revealed that females are the choosy sex in most species. This is mainly based on one assumption, the evolution of anisogamy, where males produce many small (cheap) gametes and females less but larger (expensive) gametes. Thus, females invest more energy in the production of eggs than males invest in the production of sperm (Trivers 1972). In consequence, reproduction is more costly to females and they should choose the ‘fittest’ male to mate with. This includes those with the best possible genes to improve her offspring’s fitness and/or those who can provide vital resources (e.g. territory, nesting place, food, parental care) to increase offspring survivability and attractiveness, thereby increasing the female’s personal fitness (Fisher 1958; Hedrick 1988; Møller and Alatalo 1999). Here, choosiness is defined as an individual’s active effort to invest energy and time to assess potential mates, whereas preference is defined as an intrinsic, passive attractiveness towards specific traits of the opposite sex (Jennions and Petrie 1997; Cotton et al. 2006). However, female preferences can be overridden by dominant intrasexual competition (Qvarnström and Forsgren 1998; Härdling and Kokko 2005; Formica et al. 2016). Preferences can enhance the evolution of different mating strategies and tactics to increase reproductive output with behavioral plasticity; depending on sex, age, physiological state or operational sex ratio (Parker 1982; Gross 1996).

Nevertheless, newer studies suggest that males can be choosy too, if mate availability is high and simultaneous sampling possible (Barry and Kokko 2010), if there is variation in female quality/fecundity (Krupa 1995; Johnstone et al. 1996), and if the benefits of choosing between females is higher then the costs associated with assessing females (Edward and Chapmann 2011, and references therein). Some prerequisites are the presence of males’ ability to detect differences and a preference for particular female traits. Body size can be such a trait, i.e. indicating longevity based on good genes which could be heritable (Kokko and Lindström 1996; Møller and Alatalo 1999). However, body size usually is based on a variety of genes and environmental processes, but might simply indicate higher fecundity (Peters 1986; Shine 1988; Nali et al. 2014). Mating with a larger female thus may increase a male’s individual fitness. A male’s choice however, should not only be based on such trivial correlation, it will be impacted by trade-offs concerning its mating chances, and thus individual males indeed may follow very different strategies to access females. Some examples of male tactics are satellite males, usually being smaller than their competitors (Arak 1983; Halliday and Tejedo 1995), mate-guarding (Parker 1974), prudent mate choice (Fawcett and Johnstone 2003; Härdling and Kokko 2005), clutch piracy (Vieites et al. 2004) or even functional necrophilia (Izzo et al. 2012).

Mating systems in amphibians are diverse, and apart from environmental parameters, mostly depend on female availability over time (Wells 2007). In frog and toad species (anurans) with long breeding periods (prolonged breeders) female mate choice seems to be the rule (Wells 1977). At any given time, a few females actively choose among many calling males, often based on call characteristics (Toledo et al. 2015; Ryan et al. 2019), the quality of defended territories, or the availability of other resources to judge the males (Howard 1978; Kirkpatrick and Ryan 1991; Kokko and Jennions 2008; da Rocha et al. 2018). In lek-breeding anurans, the males aggregate in displaying arenas that do not contain any resources required by females. Females visiting these arenas ‘sample’ several males and choose a male to mate with (Bourne 1992). In lek-mating systems the operational sex ratio is highly skewed towards males and individual males are not able to monopolize females, leading to higher intrasexual competition (Emlen and Oring 1977). In contrast, in species with a short breeding period (explosive breeders) males are actively searching for mates and engage in direct male-male competition over the arriving females. Explosive breeding is characterized by an almost equal operational sex ratio, synchronized receptiveness of females and low sexual selection (Emlen and Oring 1977). In theory all males are able to mate and reproduce, but larger/more dominant males have an advantage to access and dominate receptive females during scramble competition leading to a variation in male mating success (Berven 1981; Olson et al. 1986; Höglund 1989; Vagi and Hettyey 2016).

Therefore, some males are considered to sexually dominate the females in explosive breeding systems, leaving little room for male and female mate choice if the cost for mate sampling are too high (Dechaume-Moncharmont et al. 2016). Nevertheless, male mate preferences could have evolved in explosive breeders, because female fecundity highly dependents on female body size in most anuran species (Krupa 1995; Nali et al. 2014). Simultaneous sampling of preferred females might be particular possible during the peak mating time because female availability should then be highest (Arntzen 1999; Barry and Kokko 2006). All males should prefer larger females to increase their own fitness according to adaptation theory, although preferences could be obscured by high intrasexual competition. On the other hand, costs associated with mate choice depend on male density and the frequency of different mating tactics within a breeding aggregation (Arak 1983; Höglund and Robertson 1988), as well as for instance male’s individual predation risk (Magnhagen 1991; Bernal et al. 2007), all factors which may vary already during a short breeding season (Olson et al. 1986; Vojar et al. 2015).

In this study, we investigate the mating preference of the European Common Frog (*Rana temporaria*) because it is an excellent example of an explosive breeder with male-male competition. Although former studies suggest a lack of male mate preferences in this species (Elmberg 1991), we observed non-random mating by body size and found indications of male mate preference and different mating tactics in former experiments (Dittrich et al. 2018). Larger females were paired more frequently than smaller ones and smaller sized males showed a different mating tactic to get access to females (Dittrich et al. 2018). Here, we hypothesize that all males will prefer larger females independent of their own body size, when intrasexual competition is absent and males are presented to differently sized females. Additionally, we predict small males to be faster in attempting a female to increase their chances to keep an exclusive access to the female during scramble competition.

## Methods

### Study area and species

The European Common Frog, *Rana temporaria* Linnaeus, 1758, is an explosive breeder that forms dense breeding aggregations in early spring (Gollmann et al. 2014). The males engage in scramble competition over temporally available receptive females (Savage 1961). Here, larger males have an advantage in direct combat (Arak 1983), and small males may apply a different mating tactic by being faster in amplexing females (Dittrich et al. 2018). Mostly, sexual size dimorphisms exists, were females are larger in body size than males (Geisselmann et al. 1971; Elmberg 1987), but this can vary between populations (Vojar et al. 2015; Dittrich et al. 2018).

We carried out fieldwork in southern Germany, near the village of Fabrikschleichach in Lower Franconia, Bavaria (49.924 N, 10.555 E). This area comprises roughly 140 ponds, of which *R. temporaria* uses between 35 – 40 ponds for reproduction annually. In 2019 we fenced the four ponds with the largest known breeding aggregations for the entire reproductive period (14^th^ to 28^th^ March). The fence consisted of plastic gauze (mesh size 2 mm, approx. height 60 cm) stretched between wooden poles and was monitored twice a day (morning and evening).

We collected individuals that sat at the fence or were on their way to the breeding pond, preferably collecting singles to minimize differences in reproductive status. Amplected females could potentially be affected by the application of amplexin, which was found in gland tissue under the nuptial pads of male *R. temporaria.* This is a protein similar to the plethodontid modulating factor, a pheromone that influences courtship duration in salamanders (Willaert et al. 2013). So far, it is unknown if male *R. temporaria* are able to detect differences in the female’s reproductive status (Thomas 2011). We thus always took note if individuals were encountered as singles or in pairs and tested if the former status influences preference behavior of males.

All individuals were sexed *in-situ* (males show characteristic dark nuptial pads during the reproductive period). We measured snout-vent length (SVL in mm) using a caliper (to the closest 0.5 mm), and mass using a spring scale (1 – 100 g, 1 g increments). For transport, we placed each individual singly in an opaque, 1 L volume plastic bucket with lid, which contained leaf litter to hide and a thin layer of water to prevent desiccation. The animals were kept in these buckets in the barn of the ecological field station in Fabrikschleichach (temperatures only marginally higher than at the breeding sites) until the start of the behavioral experiments, which started after a maximum of 12 hours after collection. Although this handling could cause stress, studies in Cane Toads, *Rhinella marina,* showed that stress levels will decrease after 8 hours and with low temperatures (Narayan et al. 2012a, 2012b). All individuals were released at their respective capture locations after completion of the experiments.

### Behavioral experiments

We tested the hypothesis that males prefer the largest female in the absence of intrasexual competition with a mate choice test, by placing two females of different body sizes in the same container with a single small/large male. The size difference between females in each trial exceeded 10 mm, with small females SVL being below 70 mm (n = 48, range: 48 – 70 mm, mean ± SD: 63.0 ± 5.7 mm), and large females SVL over 71 mm (n = 48, range: 71 – 89 mm, mean ± SD: 77.3 ± 4.4 mm). In the containers, either a small male (n = 23, SVL range: 56 – 70 mm, mean ± SD: 63.8 ± 4.5 mm) or a large male (n = 25, SVL range: 71 – 89 mm, mean ± SD: 76.6 ± 5.5 mm) were introduced. The allocation of individuals in the experiment was random, except the premisses of a minimum of 10 mm size difference between the females. The experiments were conducted in plastic containers (40 x 60 x 40 cm), filled with 10 L of rainwater (5 cm water depth). The species can form amplexus on the migration towards the pond in the terrestrial habitat, as well as within the aquatic habitat in the pond. The 5 cm water depth are imitating the edges of the pond were clutches are usually laid. Before starting the experiment, a non-transparent plastic sheet separated the male from both females. We let the animals acclimatize in the container for 10 min, then removed the plastic sheet and started the experiment. A web camera (Logitech C920) placed at 1.5 m height above the plastic containers recorded each experiment for one hour, even if amplexus was formed earlier.

Before starting a new experimental run, we cleaned the respective container and changed the water completely to minimize the risk of potential effects from residual chemical cues. Each animal was tested only once. If successful amplexus did not occur within the one hour experimental time, the trial was terminated. In none of the experiments spawning occurred.

We defined several variables that were recorded and analyzed: when and towards which female the male attempted to clasp first, the number of successful and failed clasping attempts on each female, and with which female successful amplexus occurred at the end of the experiment. The term attempt is defined by actively approaching a female and trying to clasp her, it does not apply if animals are randomly bumping into each other.

We used χ^2^ tests to analyze male mate preferences. First, we tested if males approaches towards the small/large female were random and second, if a successful amplexus with a small/large female was random. Additionally, we tested size-assortment in the pairs with a Pearson correlation. To account for the experimental design – males can only choose from two females, thereby limiting choice – we run a bootstrapped null-model with 1000 iterations of random choice for one of the females in the experimental pairs. For those random pairs we calculated the Pearson correlation coefficients and plotted them in a histogram. We than calculated the z-score for the Pearson correlation of the observed pairs with the mean and standard deviation of the bootstrapped null-model. The same approach was used to investigate the body size difference between the approached females and successfully formed pairs. The size difference between the male and a randomly chosen females in the experimental run was calculated with 1000 iterations and plotted as the distribution of the null-model. We calculated the mean size difference for attempted pairs and for pairs in amplexus. The observed data were compared to the mean and standard deviation of the null-model. The deviation from the null-model was significant when the z-score of observed values was above 1.96. For all analysis and graphs we used the R statistical environment (R Core Team 2020, version 3.6.3). We used the packages effsize (Torchiano 2019) to calculate Cohen’s D, ggplot2 (Wickham 2009) for drawing graphs and plyr to count number of occurrences (Wickham 2011).

## Results

First, we tested if female and male status (caught as a single or in amplexus) had an influence on amplexus during the experimental runs. The proportion of females being in amplexus was not significantly different between females caught as single or paired (χ^2^ = 0.05, df = 1, *p* = 0.83). The same was observed for the males (χ^2^ = 0.17, df = 1, *p* = 0.68). Therefore, we pooled the data.

From our 48 experiments, 32 ended in the formation of pairs; 16 experiments were terminated without formation of pairs after one hour. This high number of experiments without amplexus is due to three reasons. First, a high failure rate in grabbing a female. In total, males approached females 255 times and failed to grab them in 179 cases (70.2%). In five experiments, males failed to grasp a female because she swam or jumped away. Second, in nine cases, males were successful in amplexing a female, but females showed avoidance behaviors and escaped the male grip (unp. data). Third, in two cases the males did not even tried to grab a female and showed no interest at all.

The proportion of females in amplexus was not significantly different between small (n = 13) and large females (n = 19, binomial test, *p* = 0.38). The proportion of males in amplexus was not significantly different between small (n = 15) and large males (n = 17, binomial test, *p* = 0.49). There was no indication for a preference of any female body size category by either small or large males = 0.18, df = 1, *p* = 0.67; Table 1). Additionally, during their first approach large and small males did not show any preference for a specific female body size category and tried to clasp large and small females almost equally (χ^2^ = 0.01, df = 1, *p* = 0.96; Table 1). A total of 31 males approached both females during the experiment (65%), 15 approached only one of the females (31%), and two males did not show any interest in any female (4%). The number of total attempts per male, which could be a proxy for the effort invested, were not influenced by being paired before the experiment (Welsh two-sample t-test, t = −0.41, *p* = 0.69), nor did it correlate with male body size (Pearson correlation, r = 0.13, df = 46, *p* = 0.37).

**Table 1.**
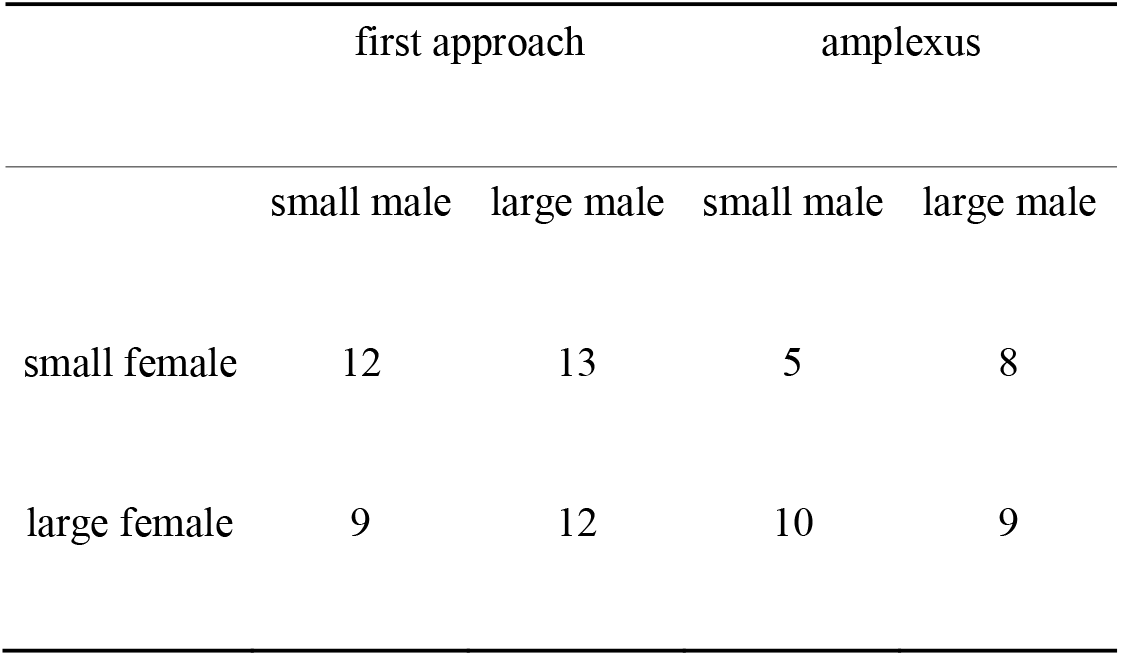
Counts of small and large males first approach towards small and large females and counts of males in amplexus with small or large females

|  | first approach |  | amplexus |  |
| --- | --- | --- | --- | --- |
|  | small male | large male | small male | large male |
| small female | 12 | 13 | 5 | 8 |
| large female | 9 | 12 | 10 | 9 |

The only difference between small and large males, was the time till they attempted a female. Small males were faster in approaching a female in the first attempt (n = 21, mean± SD = 6.83 ± 6.73 min), compared to large males (n = 25, mean ± SD = 13.62 ± 13.3 min, Welsh two-sample t-test, t = 2.24, *p* = 0.032, Fig. 1) with a medium effect size (Cohens D = 0.63).

**Fig. 1.**
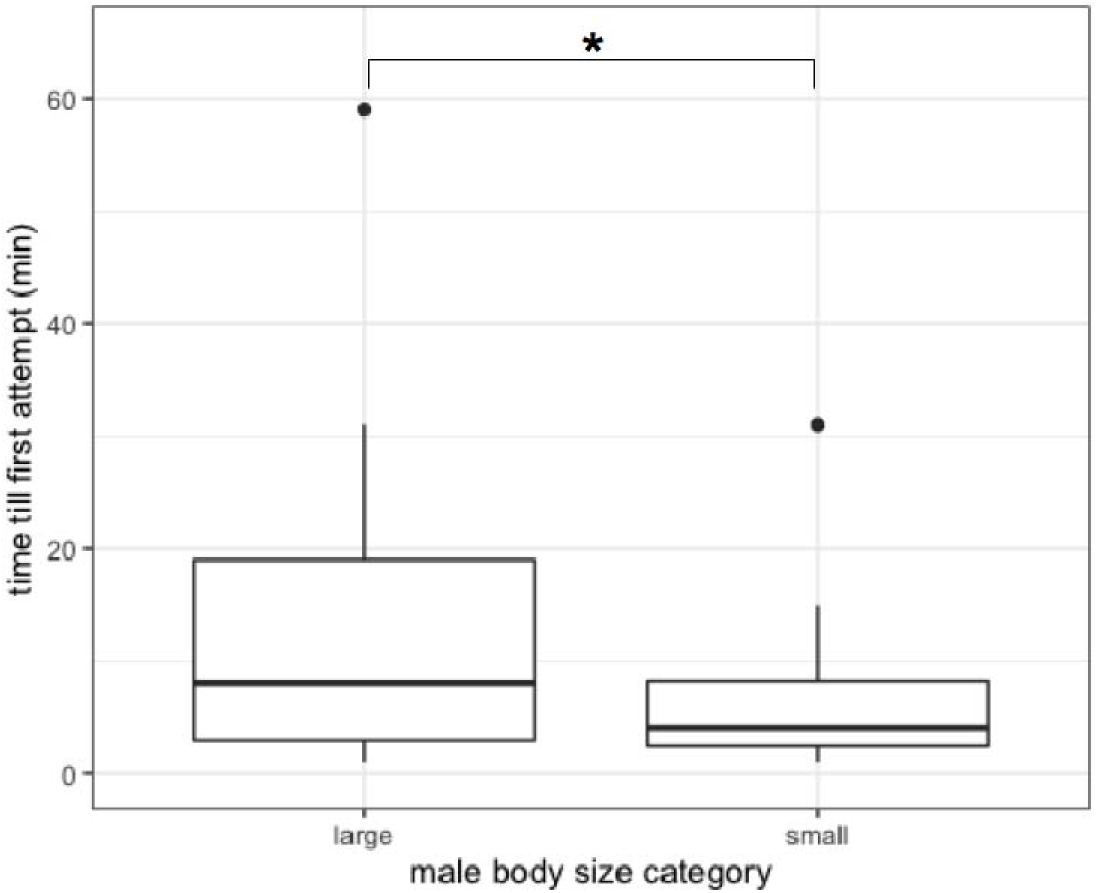
Time till first attempt of a male *Rana temporaria* to clasp a female, measured in minutes for both male body size categories. Smaller males were significantly faster in attempting a female (Welsh two-sample t-test, *p* < 0.05, depicted with asterisk). The boxplots show median (dark line), 25–75% quartile (box), non outlier range (vertical line) and outliers (black dots).

For the following analyses we did not use the size categories but took the original SVL values in mm. We observed a correlation of body sizes for final pairs in amplexus (Pearson correlation, r = 0.41, df = 30, *p* = 0.02), and also towards the first females that were attempted (Pearson correlation, r = 0.34, df = 44, *p* = 0.02). We compared these findings with a simulation of random pairings with a bootstrapped null-model. Although, allocation to the experimental pairs was random, the null-model showed a mean correlation coefficient of 0.3 and a standard deviation of 0.1. Therefore, our observed correlations were not statistically different from the random null-model (Fig. 2).

**Fig. 2.**
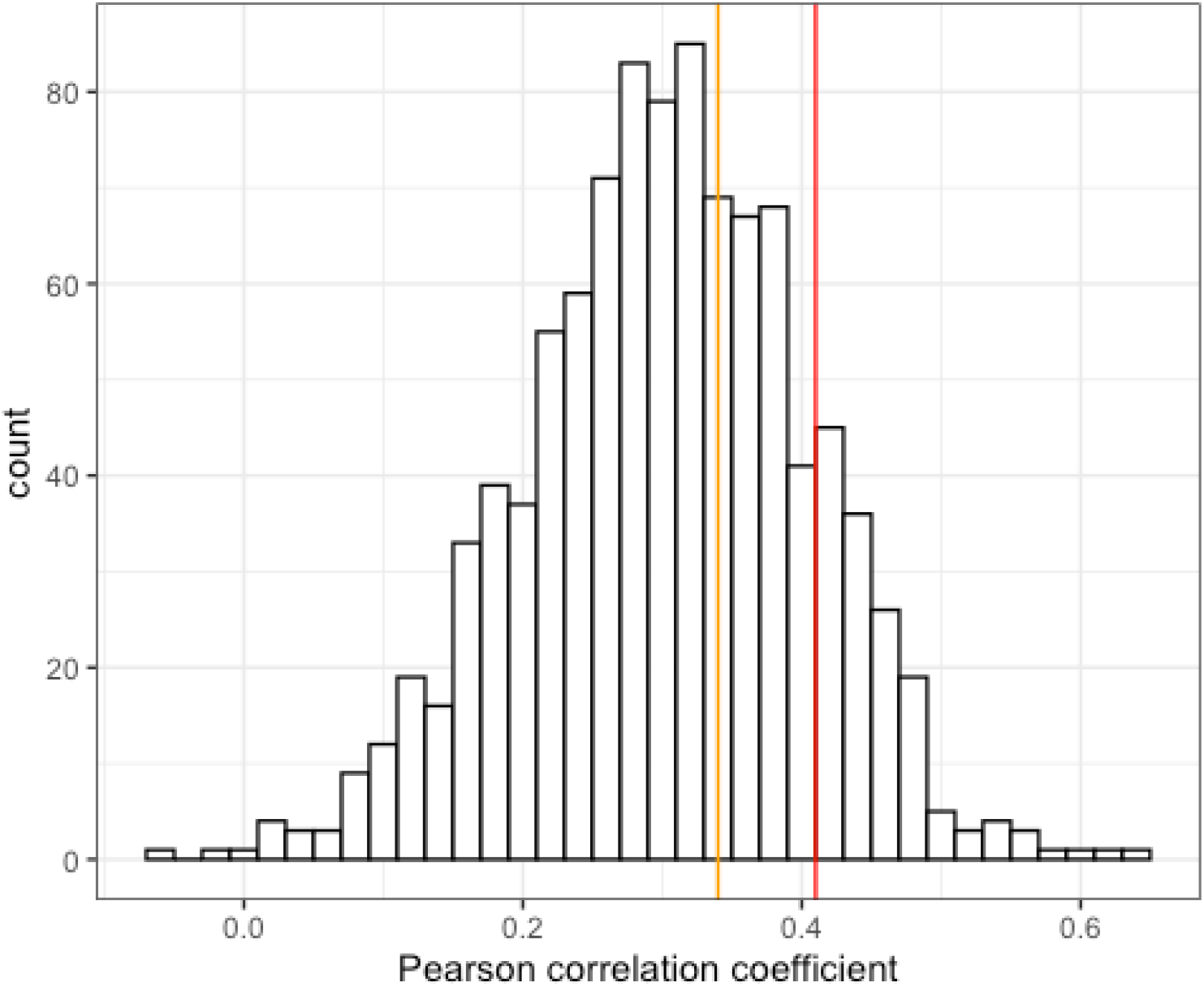
Null-model distribution of the Pearson correlation coefficient if pairing would have been random in the 48 *Rana temporaria* pairs (1000 iterations of picking one of the presented females within an experiment randomly). The orange line depicts the observed Pearson correlation coefficient of males and the first attempted female (z-score = 0.4), the red line depicts the observed Pearson correlation coefficient of the pairs in amplexus (z-score = 1.1). Both z-scores are within the boundaries of the 1.96 standard deviation from the mean value and thus random. Histogram bandwidth = 0.02.

Nevertheless, it seemed that body size difference between males and females played a pivotal role in pair formation and staying amplexed. Pairs in amplexus had a negative size difference to each other which means that the female was on average larger than the male (mean ± SD = −2.22 ± 8.42 mm). Contrary, the size difference between the male and the female that was approached first was positive, which means that the female was on average smaller than the male (mean ± SD = 1.72 ± 9.99 mm). The distribution of a bootstrapped null-model had a mean of 0.26 mm size difference between pairs and a standard deviation of 1.03 mm, if pairs were formed randomly. Therefore, the size difference of pairs in amplexus was non-random (z-score = −2.41, Fig. 3).

**Fig. 3.**
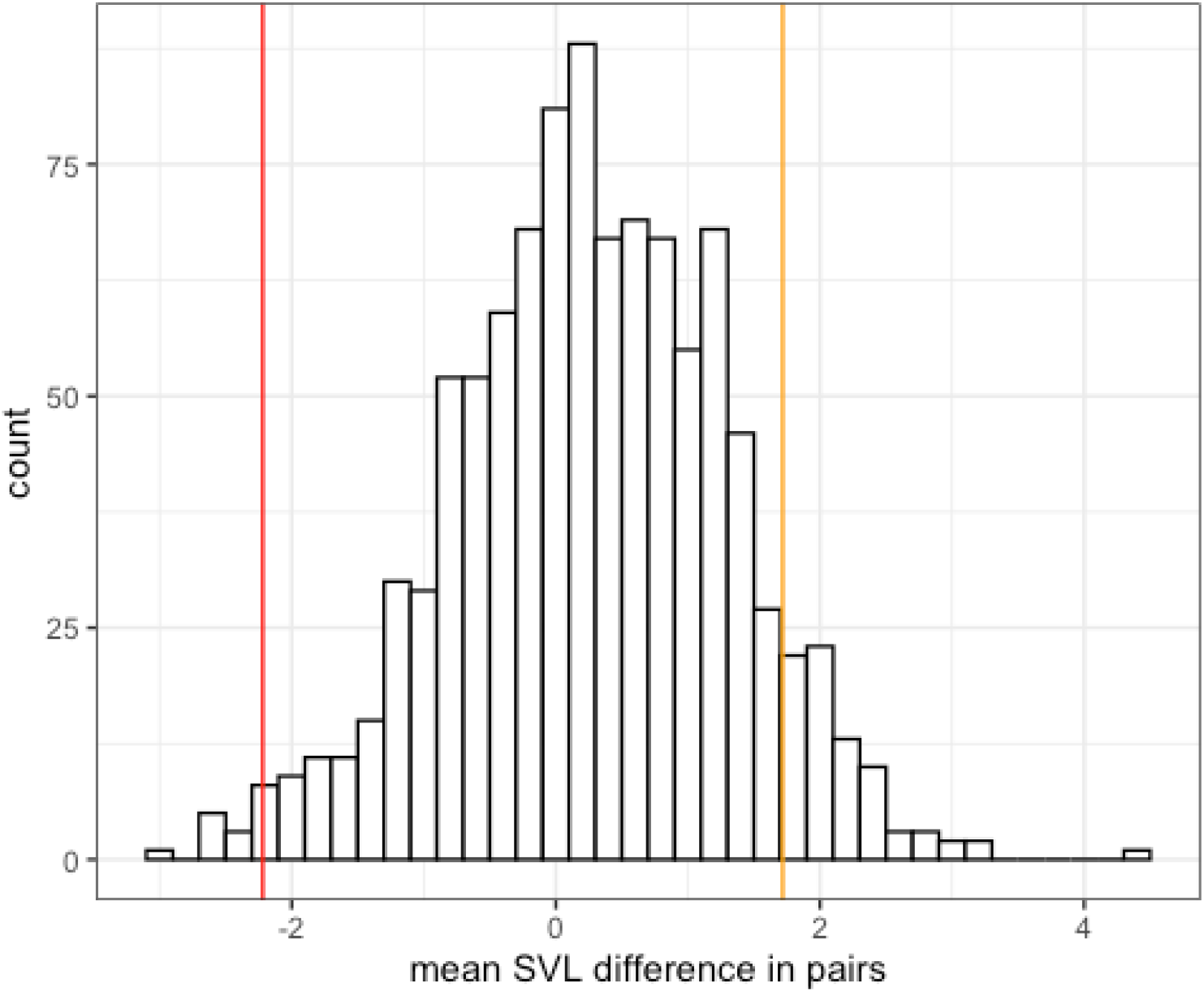
Null-model distribution of mean SVL difference in mm if pairing would be random in the 48 *Rana temporaria* pairs (1000 iterations of picking one of the presented females within an experiment). The orange line depicts the observed mean SVL difference of males and the first attempted female (z-score = 1.4), the red line depicts the observed mean SVL difference for the pairs in amplexus (z-score = −2.4). The z-score of the pairs in amplexus is larger than the 1.96 standard deviation from the mean value and thus non-random. Histogram bandwidth = 0.2 mm.

## Discussion

Based on previous observations (Dittrich et al. 2018), we expected male European Common Frogs (*Rana temporaria*) to prefer larger, more fecund females for mating. Contrary to our expectations *R. temporaria* males did not prefer any female based on body size and seem to randomly attempt females. However, as expected, small males tried to clasp females faster, which indicates a different mating tactic to increase mating chances during scramble competition. In general, the size difference between formed pairs was on average negative and significantly larger than the expected values from a bootstrapped null-model. Hence, females in successfully formed pairs were larger than the males. Because size difference between first attempted females and males was positive in our experiments, i.e. males were larger than the attempted female, a selective mechanisms other than male competition and male mate preferences seems to be responsible for this pattern.

One prerequisite for mate choice to evolve would be existing preference for specific traits, which we could not detect for female body size. This confirms Elmberg (1991), who shows that there is no male mate preference for larger body size in *R. temporaria*. However, body size alone should not be a trait under sexual selection in anurans anyhow, because it is age and resource dependent (Halliday and Verrell 1988; Lodé et al. 2004) and cannot be considered a true sexual secondary trait that provides an honest signal concerning mate quality, because heritability should be low. We did not find direct tests for heritability of body size in anurans. Although, larger body size could be an indirect indicator for good genes (Kokko and Lindström 1996; Møller and Alatalo 1999), as larger females could be better survivors and foragers. Additionally, some individuals could have an intrinsic higher juvenile growth rate which could be passed on to their progeny (Halliday and Verrell 1988), making them more attractive for mates (high juvenile growth enhances survival probabilities). Nevertheless, evidence of heritability and reliability of these traits are scarce (Hettyey et al. 2010). However, female body size is highly correlated with fecundity (Nalie et al. 2014; Dittrich et al. 2018) and this alone would be an honest signal for higher reproductive output.

In most species females grow larger than males, but males reach maturity earlier (Monnet and Cherry 2002). Indeed, within amplected pairs the females had been on average larger than the males. Although, one could argue that females are in general larger than males, in our experiments larger males approached smaller females first and the null-model indicated that pairs should be of similar size, if pair formation would be random. The explanation why this pattern is seen in the pairs which finally formed successfully, could be mechanical and independent from male preferences. If females are smaller than the amplexing male, males’ may not be able to hold them tight enough to maintain amplexus. In Cane Toads (*Rhinella marina*) it was shown that males with shorter arms could cling better to females compared to males with longer arms, the latter being replaced more often as they could not hold the females properly (Clarke et al. 2019). However, in the Woodfrog *(Rana sylvatica),* a species morphologically and ecologically very similar to the European Common Frog, longer arm length was beneficial to maintain amplexus (Howard and Kluge 1985). As we excluded male competition, the observed pattern should be due to other reasons than scramble competition and alternative mating tactics, mainly due to female avoidance behavior (unp. data).

However, mate preferences are often based on multiple cues like body size plus coloration, call characteristics, chemical cues and/or genetic incompatibility (Engeler and Reyer 2001; Taylor et al. 2007; Willaert et al. 2013; Starnberger et al. 2014; Bossuyt et al. 2019), which could not be tested in this study.

A very surprising result of our study revealed that 70% of all male attempts to clasp a female failed. A high failure rate when attempting to clasp a female should favor non-choosiness in males, due to the evolutionary cost of failing to reproduce in a given season (Krupa 1995; Fawcett and Johnstone 2003; Dechaume-Moncharmont et al. 2016). Time constraints can be an important factor in the evolution of mate choice strategies (Sullivan 1994), and time is a limiting factor in explosive breeding species such as the European Common Frog. The breeding duration highly depends on weather conditions and the time frame varies considerably between populations and locations, from a couple of days to more than two weeks (Dittrich et al. 2018). The assumptions on the evolution of mate choice and the consequences of operational sex ratio between explosive *versus* prolonged breeding are just ends of a continuum (Wells 1977), and mating strategies could vary accordingly with the length of the breeding period. Indeed explosive breeders have been reported to exhibit high intraspecific variability of mating patterns between and within population, i.e. with respect to large male advantage, different mating tactics and size-assortment, due to high variability in mate availability, as well as intrasexual competition and environmental conditions (Olson et al. 1986; Vieites et al. 2004; Vojar et al. 2015; Dittrich et al. 2018). Male mate choice as well as its absence was found in various studies addressing the same species, e.g. male mate choice detected: *Bufo bufo* (Arntzen 1999), *Rana sylvatica* (Berven 1991), no male mate choice detected: *B. bufo* (Höglund and Robertson 1987; Marco and Lizana 2002), *R. sylvatica* (Howard and Kluge 1995; Swierk and Langkilde 2019). Therefore the system of explosive, scrambling breeders seems to be context dependent which makes generalization almost impossible.

Male European Common Frogs are usually the first to arrive at breeding ponds and stay longer than females (Savage 1961; Geisselmann et al. 1971), and male mating success is positively correlated with the amount of time spent at breeding sites (Woodward 1982). However, if males spend too much time with selecting particular females or amplexing/defending a non-receptive female, males’ chances to reproduce in a given year decrease over time. Thus, males should minimize female selection time in order to increase the probability to reproduce in a given year and in the best case, reproduce more than once. That small males are faster in amplexing a female was shown before (Dittrich et al. 2018), and here we show that they are also faster in attempting to clasp a female. Small males are less competitive during scramble, but in our experiment, we excluded intraspecific competition, and thus this behavior appears to be intrinsic to smaller males and independent from other males presences. Therefore, this tactic seems to follow an individual, condition dependent strategy based on male body size (Gross 1996; Brockmann 2001). This mating system implies that until a specific body size is reached, the tactic of fast clasping is probably more successful than participating in scrambling and that a switch point exists were the reverse is true for larger body size. It would be interesting to test this assumption with the same males at different size and age. However, it should be always beneficial to get access to a female as soon as possible and to stay with the female until spawning occurs, independent of the males’ body size. Hence, the size dependent ‘speed’-hypothesis needs to be tested, including frequency and density dependent approaches, to examine reproductive success of alternative tactics (Brockmann 2001).

In conclusion, we could show that male European Common Frogs do not prefer larger females and seem to mate randomly. Smaller sized males seem to follow an individual, condition dependent strategy to get access to females compared to their larger competitors that benefit from scramble competition. However, there is a non-random mating pattern of males being in amplexus with larger females, which indicates that there are selective mechanisms that do not depend on male mate prefernces or male-male competition. Probably there are other selective forces which shape the observed mating pattern, possibly due to sexual conflict (female avoidance behavior) or female preferences.

## Ethics approval

The government from Lower Franconia issued research permits (55.2 DMS 2532-2-316) and the Bavarian state forestry department provided access the forest ponds. All animal behavioral experiments followed the guidelines provided by ASAB.

## Availability of data and material

The datasets generated and analyzed during the current study, including the R script, are available at the Museum für Naturkunde repository, doi: 10.7479/zpnq-pj77

## Acknowledgments

We thank the government from Lower Franconia for research permits and the Bavarian state forestry department in Ebrach for access permission to the forest ponds. The staff of the ecological field station, University of Würzburg, provided access to crucial infrastructure, which is highly appreciated. We thank Olga Ritz for her help analyzing the video material and Sami Asad for proof reading concerning English grammar and style.

